# Detection of an unstable minor QTL linked to fire blight resistance on linkage group 16 of *Malus fusca*

**DOI:** 10.1101/2021.06.04.447050

**Authors:** Ofere Francis Emeriewen, Klaus Richter, Annette Wensing, Mickael Malnoy, Andreas Peil

## Abstract

**Objective:** *Erwinia amylovora* causes fire blight disease in *Malus*. A strong resistance QTL (*Mfu10*) was previously detected on linkage group 10 of *Malus fusca* accession MAL0045, using several strains of the bacterium. As no strain capable of breaking the resistance of MAL0045 has been found, it was hypothesized that a second resistance factor contributes to the fire blight resistance of MAL0045. However, to date, no minor locus has been detected with previously published strains of the bacterium. We detected a minor QTL only on a subset of a population following inoculation with strain Ea1038, which heterologously expresses an effector in a derivative of isolate Ea3049. Two genetic maps of MAL0045, one scarce, the other dense with markers, were used for QTL analyses.

**Results:** *Mfu10* was detected on LG10 with Ea1038, as was previously with Ea3049. Although no other QTLs of significant LOD was previously detected in other linkage groups with Ea3049, a QTL of significant LOD was detected on LG16 (*Mfu16*) after inoculation of a subset of 76 individuals with Ea1038, but only using the dense genetic map. *Mfu16* improved the effect of *Mfu10*. However, when the number of individuals inoculated with Ea1038 was increased to 121, *Mfu16* was no longer detected in the dense genetic map. We hypothesize some factors, which might be responsible for the instability of this QTL.

## Introduction

The bacterium, *Erwinia amylovora*, causes fire blight – a devastating disease of the domesticated apple (*Malus domestica* Borkh.) and related species (*Malus* spp.) [1–4]. The mechanisms by which the pathogen invades and causes disease in susceptible hosts have been extensively reviewed [2, 5]. Similarly, the molecular strategies employed by resistant hosts for the recognition of *E*. *amylovora* elicitors are well documented [2, 4, 6]. *Malus* host resistance is mostly quantitative, evidenced by the segregation of resistant and susceptible phenotypes in *Malus* populations [7–11]. Quantitative trait loci (QTLs) for fire blight resistance have been detected in apple cultivars and wild apple species accessions [reviewed in 4]. Although most apple cultivars are more susceptible to the disease, their QTLs are faster to introgress but not sufficient to provide strong resistance as those of their wild relatives [8–11], for which fire blight resistance candidate genes have been proposed [12–15], and in one instance, functionally proven [16].

A strong fire blight resistance QTL (*Mfu10*) was detected on linkage group (LG) 10 of the wild apple accession, *Malus fusca* MAL0045 [9]. The stability of *Mfu10* has been demonstrated using several strains of *E*. a*mylovora* differing in virulence/aggressiveness [17–19]. In particular, the highly aggressive Canadian strain, Ea3049, and the mutant strain ZYRKD3-1, both of which break down the resistance locus of another wild *Malus* genotype, *Malus ×robusta* 5 (Mr5) [20, 21], could not breakdown *Mfu10* [17, 18]. However, unlike in Mr5, where minor QTLs were detected following inoculation with Ea3049 [22], no minor QTLs were detected in MAL0045 following inoculations with four different strains [19].

### Rationale and methods

It was previously reported that a switch from cysteine amino acid (C-allele) to serine amino acid (S-allele) in the avrRpt2_EA_ effector protein sequence of *E*. *amylovora* at position 156 is responsible for virulence and resistance breakdown in Mr5 [21] but also aggressiveness in other *Malus* host [4]. Ea3049 possesses the S-allele contributing to the high virulence/aggressiveness of this strain.

That MAL0045 itself is highly resistant to Ea3049 but the average PLL of the progeny increased to 62.4 compared to of 22.6 after inoculation with strain Ea222 [17], led us to speculate that there might be a second factor contributing to the resistance of MAL0045, and/or the SNP in the avrRpt2_EA_ effector of Ea3049 [21] might not be the only virulence factor of this strain. We inoculated the original mapping population (05210 individuals) derived from MAL0045 × ‘Idared’ cross [9], with a mutant strain of Ea3049. In the meantime, we developed a dense genetic map of MAL0045 using genotyping-by-sequencing generated SNPs incorporated with microsatellite markers (SSRs), for MAL0045-derived progenies namely: 05210 and 09260 individuals [19].

The strain used in this study, Ea1038, is a derivative of Ea3049 with the chromosomal S-allele of the *avrRpt2_EA_* effector deleted and complemented with the C-allele on an expression vector. Artificial shoot inoculation was performed on scions of up to 10 replicates of each progeny individual grafted onto rootstock M9, by cutting the youngest leaves with a pair of scissors dipped into bacterial inoculum (10^9^ cfu/ml). Disease necrosis was measured in centimeters 27 days post inoculation (dpi) and converted into percent lesion length (PLL) by dividing the necrotic shoot by the total shoot length and multiplying by 100. The preliminary QTL mapping was done with the phenotypic data of 76 inoculated plants. Subsequently, the number of phenotyped individuals could be increased to 121 individuals. These data were used for a second QTL mapping. The calculated average of PLL of all replicates of each individual was used for Kruskal-Wallis analysis and interval mapping using MapQTL 5.0 [23]. The first incomplete genetic map of MAL0045 [9, 24] and the recently developed dense map [19] served as templates for QTL analysis. This map was established using 112 individuals of the 05210 population and 36 individuals of the 09260 population, 148 individuals in total.

SAS (SAS Institute) GLIMMIX (generalized linear mixed model) analysis was performed to determine whether the effects of detected loci were significantly different. For this analysis, phenotypic values (PLL) of each progeny individual as well as their marker alleles were employed.

## Results

Of 112 individuals of 05210 population used to develop the map [19], it was only possible to phenotype 76 individuals by artificial shoot inoculation with Ea1038 in 2017. The distribution of PLLs for all 76 individuals is shown in Figure 1. For these individuals, 61.1 % and 65.7 % were the mean and median PLLs, respectively. Only five individuals recorded PLLs below 10.0 with 3.3 the lowest. Fifty-two individuals recorded PLLs over 50 with 100 recorded as the highest for twelve individuals. For the parents, whilst a PLL of 9.6 was recorded for MAL0045, 100 % was recorded for ‘Idared’.

**Figure 1.**
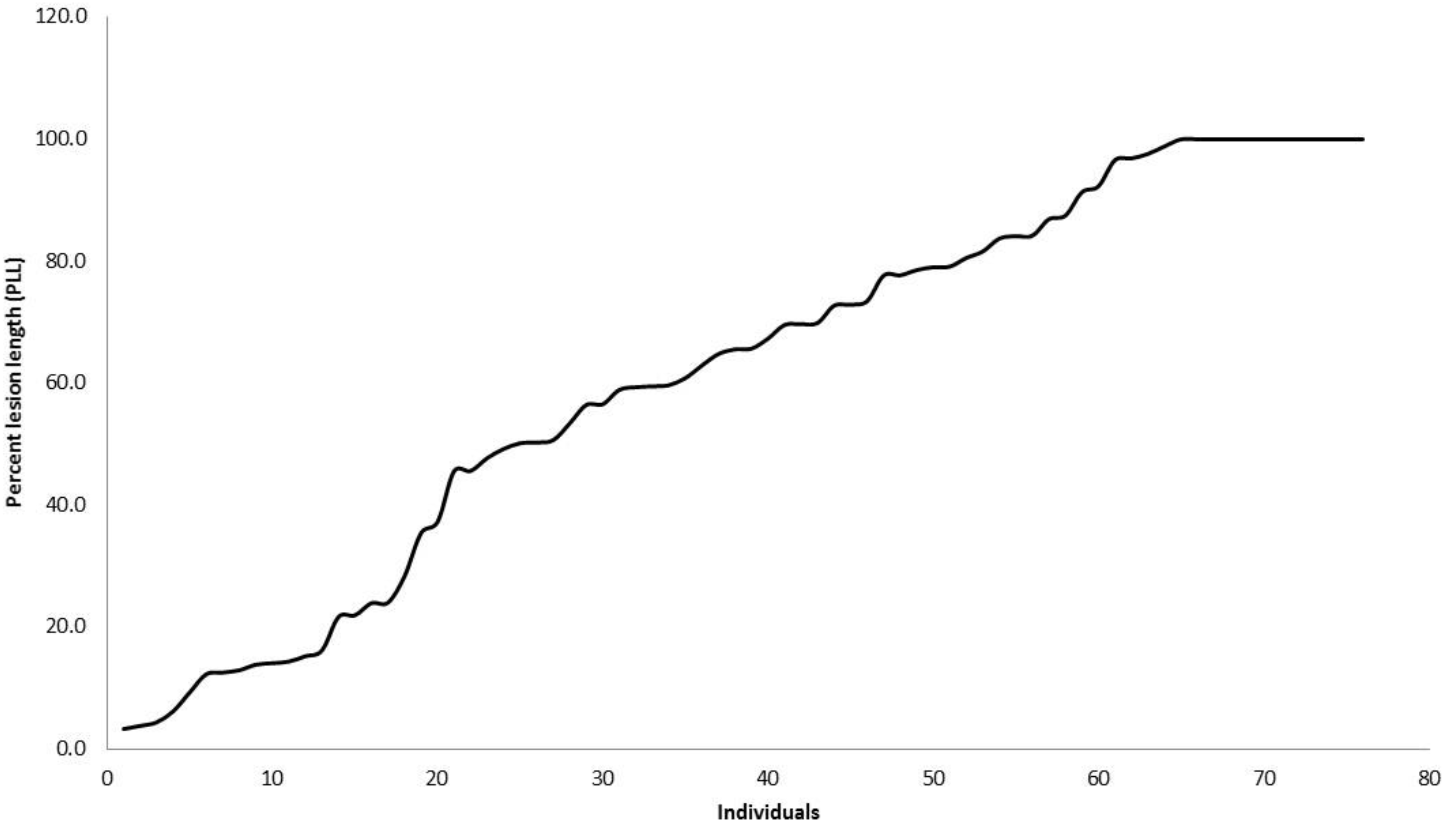
Distribution of PLL of seventy-six individuals of 05210 population phenotyped with Ea1038

The averages of all replicates for each of the 76 individuals were used for Kruskal-Wallis analysis and interval mapping with the newly developed map as template. Kruskal-Wallis analysis showed that markers on LG10 (highest *K* value = 37.2), LG16 (highest *K* = 13.0) and LG17 (highest *K*= 11.6) correlated with fire blight resistance (Table 1). The most significant correlation with resistance was observed in LG10 where individuals inheriting the resistant allele of the marker with the highest *K* value (Sca_304010_602250) possessed 43.8 % less necrosis than individuals inheriting the susceptible allele (i.e. difference between mean ll and mean lm genotypic segregation). For LG16 and LG17, the differences were 28.1 % (Sca_304000_613165) and 22.1 % (CH05b06_L4), respectively (Table 1). With the first incomplete map, only markers on LG10 showed correlation with fire blight resistance (data not shown).

**Table 1.**
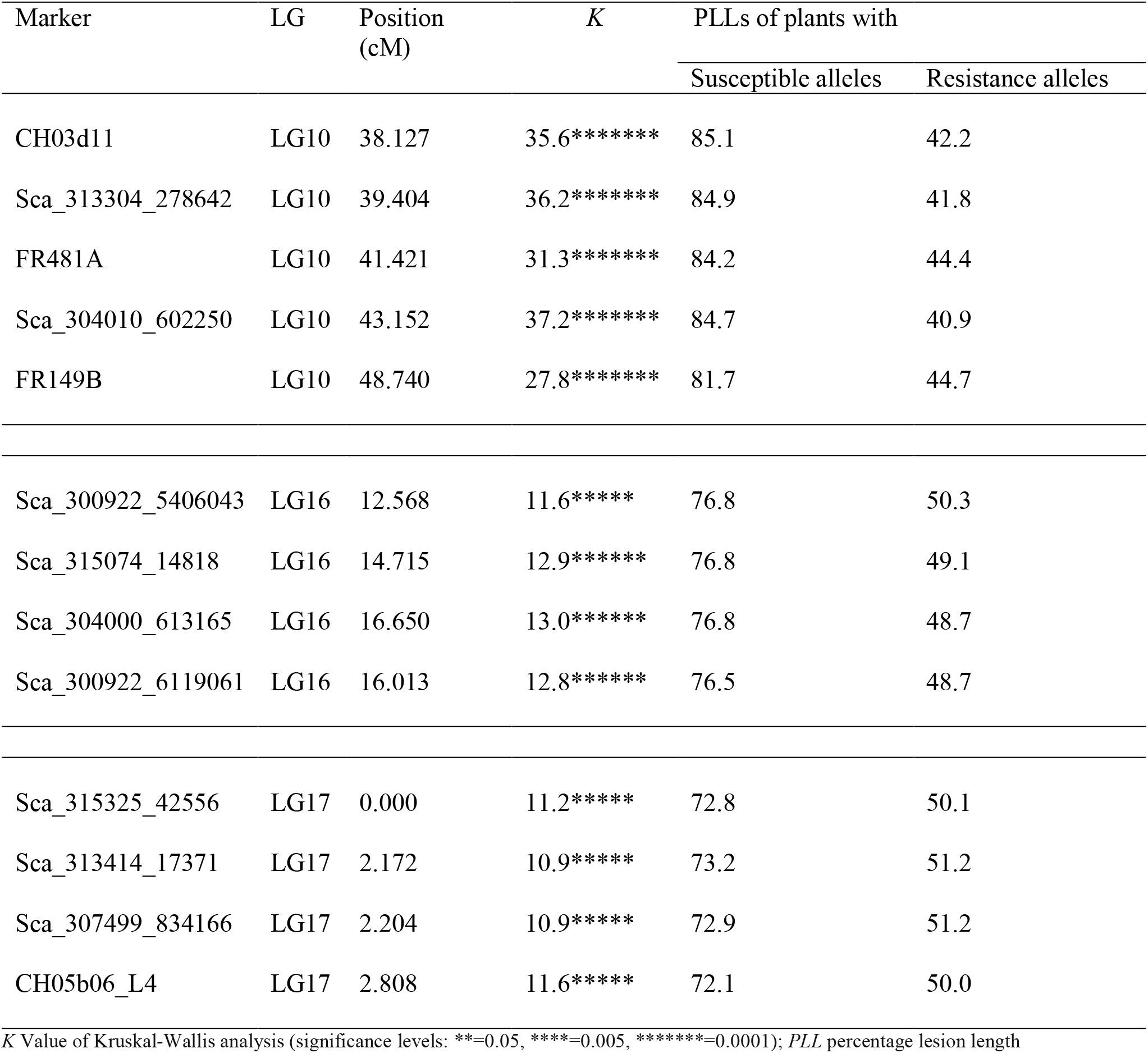
Kruskal-Wallis analysis results for some markers on the three linkage groups of the dense genetic map showing strong correlation with fire blight resistance following inoculation with Ea1038

Interval mapping with the genome wide (GW) threshold of 4.9 identified two QTLs of significant LOD scores on LG10 and LG16. No significant QTL was found on LG17. The QTL detected on LG10 is *Mfu10*, previously detected with other strains of *E. amylovora* since it is located in the same interval between CH03d11 and FR149B with markers possessing highly significant *K* values (*P* = 0.0001) (Table 1). However, the QTL on LG16 (*Mfu16*) is a novel minor QTL (Figure 2) never previously detected with any strain. *Mfu16* was not detected in the scarce map with Ea1038. Interval mapping results was in agreement with Kruskal-Wallis analysis as SNP markers on LG16 with highly significant *K* values (Table 1) possessed the highest LODs and appear underneath the QTL plot (Figure 2). The significance and interaction between *Mfu10* and *Mfu16* were determined using SAS GLIMMIX analysis. Fire blight resistance was significantly stronger when resistance alleles of both loci were present in individuals compared to when individuals possessed only *Mfu10* or *Mfu16* resistance alleles (Figure 3). Further, the resistance level of *Mfu10* alone was significantly stronger than the resistance level of *Mfu16* alone.

**Figure 2.**
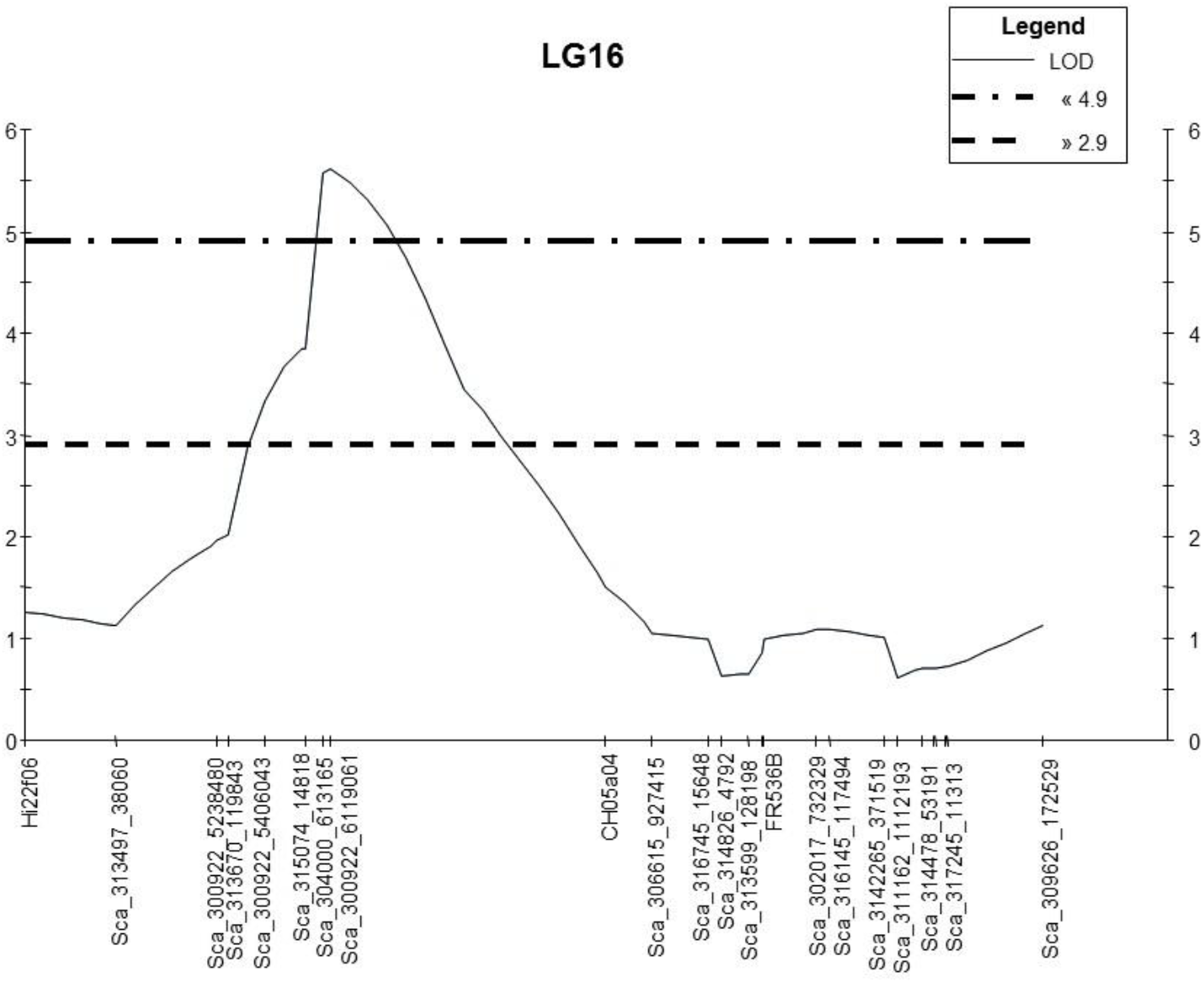
LOD plot of interval mapping for the detected QTL on LG16 showing the significance at the chromosome level threshold of 2.9 and genome wide threshold of 4.9, with the markers

**Figure 3.**
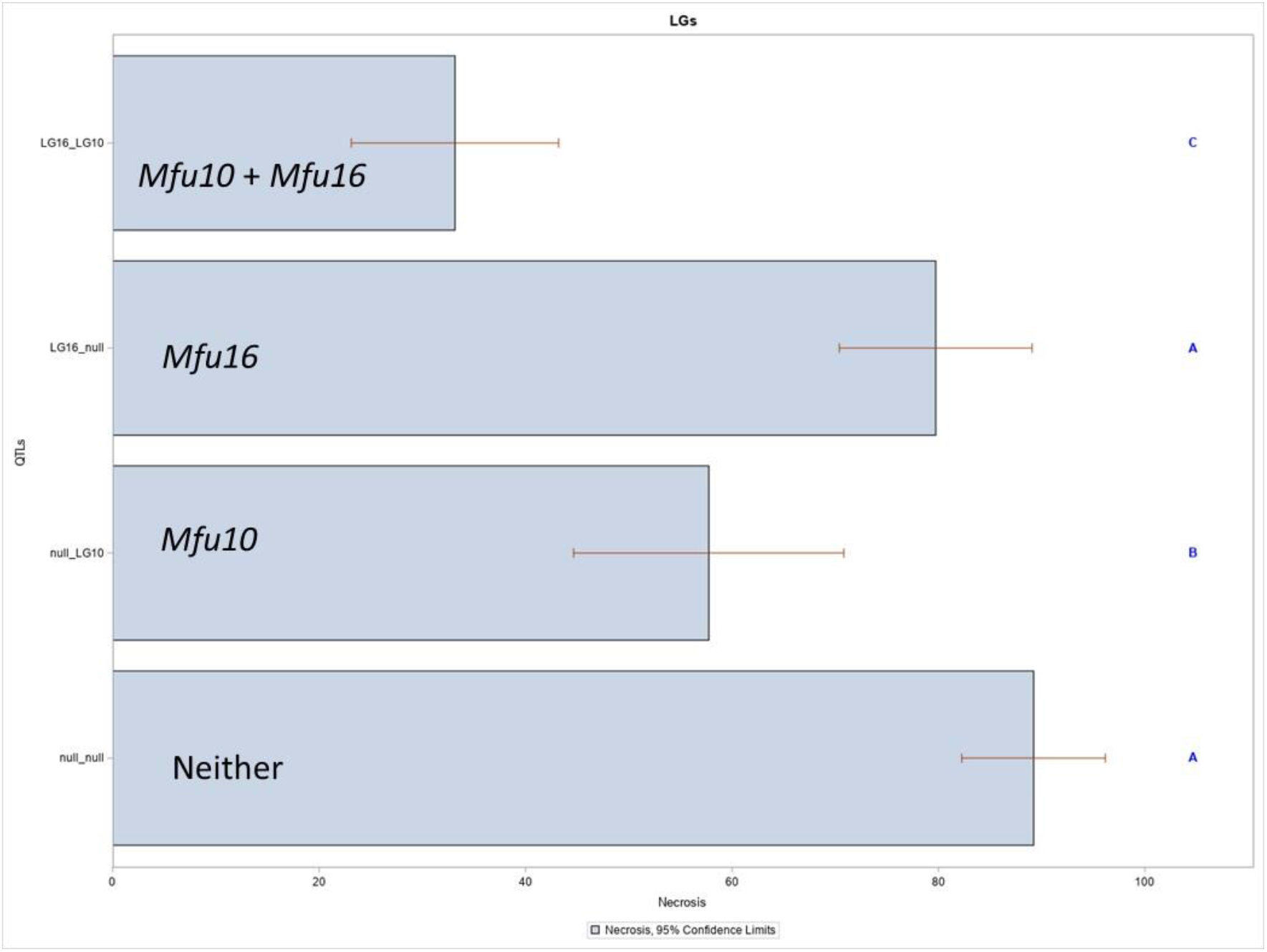
Significance and interaction between *Mfu10* and *Mfu16* determined by SAS GLIMMIX analysis

A total of 121 individuals could be inoculated with Ea1038 from 2017 to 2020. The mean PLL calculated for the 121 individuals was 55.30 with a median of 59.38. Nine individuals recorded PLLs below 10.0 with 1.09 being the lowest in only one individual. Seventy individuals recorded PLLs over 50 with 100 recorded as the highest for five individuals. QTL analysis with these data resulted in the detection of *Mfu10* (data not shown) but not *Mfu16*.

## Discussion

We study the interaction of *M. fusca* (MAL0045) and derived progeny with different strains of *E. amylovora* through artificial inoculation and QTL mapping. Through this process, *Mfu10* was first identified on LG10 using Ea222 [9, 24]. The highly virulent Canadian strain, Ea3049, affected *Mfu10* but did not overcome it, although this strain was not aggressive on MAL0045 [17, 19]. We therefore hypothesized a second putative resistance factor, possibly another locus, might be involved in the resistance of MAL0045. The failure to detect another locus was partly attributed to the fact that the first genetic map of MAL0045 was only scarce and did not represent the whole genome [9, 24]. This hypothesis was predicated on the situation in Mr5 where, although one minor QTL was detected on LG5 after inoculation with Ea3049 [20], a few more minor QTLs were detected following the development of a more saturated genetic map and inoculation with different strains [22]. The first map of MAL0045 [9, 24], developed with only the 05210 population, consists of 213 loci made up of DArT markers, a few SNPs and SSRs developed from the apple genome [25]. On the other hand, using tunable genotyping-by-sequencing technology (tGBS), thousands of *de novo* SNP markers were developed for MAL0045, of which 560 SNPs were mapped including 53 SSR markers [19]. Thus, the dense map has 400 markers more than the initial map and correctly represents the genome of MAL0045. However, no minor fire blight locus were detected with this map [19]. Since fire blight resistance is strain-dependent [21], we therefore speculated that the failure to detect a minor locus is not only dependent on the marker density of the genetic map, but also the interaction with the given *E*. *amylovora* strain used for inoculation.

Ea1038 phenotypic results showed that this mutant strain was as virulent as the wild type, Ea3049 [17] with more than half of the 76 individuals having PLLs above 50 %. The mean PLL of 61.7 with Ea1038 is similar to 62.4 recorded with Ea3049 on the 05210 population. The results are also similar to the effect of Ea3049 on another wild apple *M*. ×*arnoldiana* – MAL0004 [11], where the mean PLL recorded for 87 individuals was 65.9 %. It is interesting to note that although this mutant is complemented with the C-allele, it was still very aggressive to the individuals inoculated. This suggests that the switch from cysteine amino acid (C-allele) to serine amino acid (S-allele) at position 156 of avrRpt2_EA_ amino acid sequence [21] is not the only factor that contributes to the pathogenicity of S-allele strains. Nevertheless, we detected two QTLs of significant LODs on two different linkage groups, LG10 and LG16. It was quite clear that *Mfu10* is QTL located on LG10, however, a novel minor fire blight QTL, never previously detected with any strain was located on LG16 (*Mfu16*). We propose *Mfu16* as a minor fire blight locus albeit unstable. Although, *Mfu16* was detected only with the subset of 76 individuals, and independently did not contribute significantly to resistance levels, it positively affects *Mfu10*, as the effect of both loci is significantly stronger than *Mfu10* alone in the 05210 individuals. Minor fire blight QTLs were detected on LGs 5, 7, 11, and 14 of Mr5 [22], however, only the minor QTL on LG7 was found to contribute to resistance in addition to the major QTL on LG3 [10, 26].

Both the strain and the dense map were important factors in detecting *Mfu16* in this study. That this mutant strain and not the wild type, Ea3049, led to the detection of *Mfu16* is indicative of strong incompatible interaction between *M. fusca* and the C-allele of the avrRpt2_EA_ effector of *E. amylovora*. In addition, the failure to detect *Mfu16* in the initially developed map [9] is indicative of the important role dense genetic maps play in molecular genetics studies in *Malus* species and other plant species.

## Limitations

The failure to detect *Mfu16* following the addition of more phenotypic data is somewhat of a surprise and frankly unexplainable result, although we do not rule out the role of individuals in the map with missing phenotypic data. Some of the individuals with missing phenotypic data died off in the orchard and hence could not be phenotyped. It cannot be excluded that the exchange of genotypes during scion cutting, grafting, inoculation or measuring could be a reason, too. Nevertheless, the results are strong indications of a putative minor QTL on LG16 of MAL0045, which significantly improves the resistance of *Mfu10*. In the following years, more phenotypic evaluation of these individuals and other established crosses with MAL0045-derived progeny will help determine the usefulness and stability of *Mfu16*.

## Supplementary Information

Not applicable.

## Abbreviations

cfu/ml: colony forming units/milliliter
dpi: days post inoculations
GW: genome wide
GLIMMIX: generalized linear mixed model
LG: linkage group
LOD: logarithm of the odds
PLL: percent lesion length
QTL: quantitative trait locus
SNP: single nucleotide polymorphisms
SSRs: simple sequence repeats

## Ethics approval and consent to participate

Not applicable

## Availability of data and material

Data generated from this study are published within this article. Further materials can be provided on request from the corresponding authors, Ofere Francis Emeriewen and Andreas Peil.

## Acknowledgements

We acknowledge DFG for providing funding. We thank the orchard staff at Dresden-Pillnitz and Quedlinburg for excellent technical assistance.

## Consent for publication

We confirm that all authors read and approved this manuscript for publication

## Authors’ contributions

AP, OFE and MM for concept of the research, AP established the populations, OFE is responsible for the project and AP supervises the research; AW developed the mutant strain, KR performed the inoculations, AP and OFE performed mapping analyses, OFE and AP prepared the manuscript, and all authors read and approved the manuscript.

